# MolPhase: An Advanced Phase Separation Predictor and an Investigation of Phytobacterial Effector in Plant

**DOI:** 10.1101/2023.09.21.558813

**Authors:** Qiyu Liang, Nana Peng, Yi Xie, Nivedita Kumar, Weibo Gao, Yansong Miao

## Abstract

We introduce MolPhase (http://molphase.sbs.ntu.edu.sg/), an advanced protein phase separation (PS) prediction algorithm that improves accuracy and reliability by utilizing diverse physicochemical features and extensive experimental datasets. MolPhase applies a user-friendly interface to compare distinct biophysical features side-by-side along protein sequences. By additional comparison with structural predictions, MolPhase enables efficient predictions of new phase-separating proteins and guides hypothesis generation and experimental design. Key contributing factors underlying MolPhase include pi-pi interaction, disorder, and prion-like domain. As an example, MolPhase finds that phytobacterial type III effectors (T3Es) are highly prone to homotypic PS, which was experimentally validated *in vitro* biochemically and *in vivo* in plants, mimicking their injection and accumulation in the host during microbial infection. In addition, the phase-separation of T3Es were evolved both *in vivo* and *in vitro*, suggesting their determinative scaffolding function, though there is a difference in material properties, implying a difference in homotypic and heterotypic macromolecular condensation. Robust integration of MolPhase’s effective prediction and experimental validation exhibit the potential to evaluate and explore how biomolecule PS functions in biological systems.

## Introduction

Protein phase separation (PS) in biological systems is a complex and spatiotemporally regulated process to create functional membraneless organelles. The initiation and propagation of homotypic PS are determined by the intrinsic physicochemical properties of each biomolecule, which will also be significantly influenced by the abundance, localization, and components in living systems. When multiple macromolecular components condense, the resulting heterotypic phase separation is influenced by the complex interactions of each component. This is determined by balancing their individual biophysical characteristics and associative behaviors (Pappu *et al*, 2023). To effectively analyze the propensity for protein phase separation based on intrinsic physicochemical features is crucial. This method aids in formulating informed hypotheses by also considering their *in vivo* macromolecular environments. The initiation of protein cluster formation under subsaturation conditions is a critical transition step before evolving to high-order assembly and then the near-equilibrium phase separation, which is often shown by *in vitro* biochemical assays (Lan *et al*, 2023; Lee *et al*, 2023; Seim *et al*, 2022). The formation of initial molecular assemblies and their evolution into condensates are largely influenced by the intrinsic biophysical properties of scaffolding proteins. These processes start at a nanometer scale, making them difficult to quantify. This challenge is further compounded by the need for high spatiotemporal resolution in microscopy and sensitivity in single molecule detection. An effective prediction of homotypic PS using biophysical features of protein sequences has shown great potential to guide hypothesis generation and experimental design. Combining existing knowledge of PS candidate proteins and integrating technologies in characterizing molecular dynamics (Case *et al*, 2019; Ma *et al*, 2021; Ma *et al*, 2022; Sun *et al*, 2021), protein structure, and protein-protein interactions (Ma *et al*., 2022; Spegg *et al*, 2023; Tran *et al*, 2020) makes rational forecasting of phase separation via computational approaches feasible.

Here, we introduce MolPhase, a protein PS and molecular condensation predictor. MolPhase was trained using 606 experimental-derived PS proteins. MolPhase applies a broad set of physicochemical features and incorporates larger and more diverse experimental datasets, which showed improved accuracy and reliability among several available phase-separation-prediction algorithms (Chen *et al*, 2022; Chu *et al*, 2022; Hatos *et al*, 2022; Orlando *et al*, 2019; Saar *et al*, 2021; van Mierlo *et al*, 2021). MolPhase enables efficient analysis of extensive protein sequence datasets, facilitating the identification of novel phase-separating proteins and their functional roles. By combining with the features analysis generated by MolPhase, rationale design could be carried out to dissect the underlying PS contributing factors. Using MolPhase, we found that several physical-chemical interaction modes, including pi–pi interaction, disorder and prion-like domain, ranked as top contributing features.

To assess and validate MolPhase’s performance, we investigated bacterial type III effectors (T3Es) both *in vitro* biochemically and *in vivo* within living host cells. T3Es naturally injected into plants during microbial infection via the type III secretion system (T3SS), accumulate in the host and gradually subvert host biology based on their localization, abundance, and recognition partners in a spatiotemporal manner. Given the known phase-separating nature of T3E XopR and the prevalent highly structure disordered characteristics of T3Es, we hypothesize that many other phytobacterial T3Es could form molecular condensates on the plant’s plasma membrane and cytoplasm (Marin & Ott, 2014; Sun *et al*., 2021; Wang *et al*, 2022b). We analyzed all the T3Es from two most studied model phytobacteria *Xanthomonas campestris* pv. *campestris* (*Xcc*) 8004 and *Pseudomonas syringae* pv. *tomato* (*Pst*) DC3000. According to the prediction of MolPhase, we chose a few representative T3Es and experimentally examined their PS in living plants via cell imaging and biochemical behavior using recombinant proteins. Our findings revealed that these T3Es undergo PS both *in vitro* and *in vivo*, consistent with MolPhase analysis. However, the homotypic PS from recombinant proteins showed different material properties and dynamics than the condensates formed in living cells. It suggests a formation of heterotypic PS *in vivo* by recruiting other plant components, in which T3E might function as a scaffolder that recruits and manipulates plant molecules for subversion. Additionally, we conducted a global analysis of the *Xcc* 8004 and *Pst* DC3000 proteomes and compared them with the *Pseudomonas syringae* Type III Effector Compendium (PsyTEC), which contains 529 T3Es (Laflamme *et al*, 2020). Compared to the entire phytobacteria proteome, T3Es *Pseudomonas* species exhibit a much higher PS propensity, indicating the potential universal mechanism by which bacterial T3Es subvert host biology through their abilities in multivalent interactions and phase separation.

## Results

### Feature characterization for machine-learning predictor

To expand the pool of protein sequences that undergo phase separation (PS) as training data, we retrieved sequences from public databases: LLPSDB (Wang *et al*, 2022a), PhaSePro (Mészáros *et al*, 2020), DrLLPS (Ning *et al*, 2020), PhaSepDB (Hou *et al*, 2023) and CD-CODE (Rostam *et al*, 2023), along with additional 99 sequences curated from literature manually. For all the above-mentioned sequences, only those with experimental evidence that can undergo homotypic PS were selected to construct the final training set (Appendix Dataset 1). To remove the redundant sequences with high similarity, CD-HIT was applied to filter sequences with an identity higher than 0.9 (Fu *et al*, 2012). In total, we obtained 606 sequences for positive PS dataset (designated as POS) to study PS protein property and to model training in the subsequent step. Sinc homogenously tightly-fold structured protein will not undergo multivalent interaction based PS on their own, we selected 1367 proteins from PDB (Berman *et al*, 2000) as the negative PS dataset (designated as NEG), all training set sequences were included in Appendix Dataset 1.

To characterize the features that contribute to PS, here we dissect multiple biophysical and biochemical properties between POS and NEG (Fig 1A-L, Appendix Table 1). Compared with NEG, POS contains longer sequences (Fig 1A). The increasing in sequence length can reduce the entropic cost of confining the protein in a dense phase (Martin & Mittag, 2018). Given that PS is driven by the multivalent interaction of different domains, we explored the characteristics and discrimination of intrinsically disordered regions (IDRs) and low complexity regions (LCRs) in protein sequences from POS and NEG (Fig 1B-D). POS contains a higher percentage of IDR and LCR (Fig 1B and C). This is consistent with the enriched interacting motifs within IDR and LCR that serve as the “stickers”, which drive molecular condensation in the combination with “spacer” (Li et al., 2020)(Martin *et al*, 2020), including prion-like domain (PLD), type II polyproline helices (PPII) (Brown & Zondlo, 2012), pi interaction (including pi-pi/pi-cation interaction) (Vernon *et al*, 2018), charge-charge interaction, and the hydrophobic effect. Fig 1E-G show that POS contains a higher fraction in interactive domains. In contrast, we found that fraction of charged residues (FCR) is lower in POS (Fig 1H), and net charge per residue (NCPR) is closer to zero in POS (Fig 1I). Additionally, we introduced kappa and omega to indicate the distribution of charged residues pattern in the whole sequence. In detail, kappa value reflects the mixture of charged amino acid within the whole sequence (Das & Pappu, 2013), and omega reflects the mixture of charged residues plus proline in the sequence (Martin *et al*, 2016). Lower kappa and omega values indicate a superior mixture, whereas higher values suggest the formation of more localized blocks. It’s noteworthy that charge patterns exhibited a better mix in NEG (Fig 1J). However, when proline and charge amino acids were combined, there was no significant difference (Fig 1K). When considering FCR, NCPR, and kappa together (Fig 1H-J), it appears that proteins prone to PS generally maintain a neutral charge. Yet, they present more localized charge blocks, as indicated by kappa. The reason why omega doesn’t show a significant difference between POS and NEG (Fig 1K) remains unclear. This suggests that most proteins adapt to the physiological cellular neutral pH condition without inherent homotypic PS. Instead, they facilitate inter- and intramolecular interactions using localized charge blocks and other interactive motifs as needed. Regarding hydrophobicity, POS is more hydrophilic than NEG (Fig 1L). Interestingly, while local hydrophobic cluster/domain could enhance coacervation (Yeo *et al*, 2011) and stabilize phase separating FUS (Krainer *et al*, 2021), a lower overall ratio of hydrophobic residues is believed to maintain amino acid chains in a disordered states, potentially facilitating condensation in more liquid-like state(Dignon *et al*, 2020). This implaies that the role of hydrophobicity in guiding molecular condensation requires protein-specific and position-based consideration. In our findings, POS has a lower hydrophobicity than NEG, whereas charge blocks directly drive PS (Fig 1 E-L).

**Figure 1.**
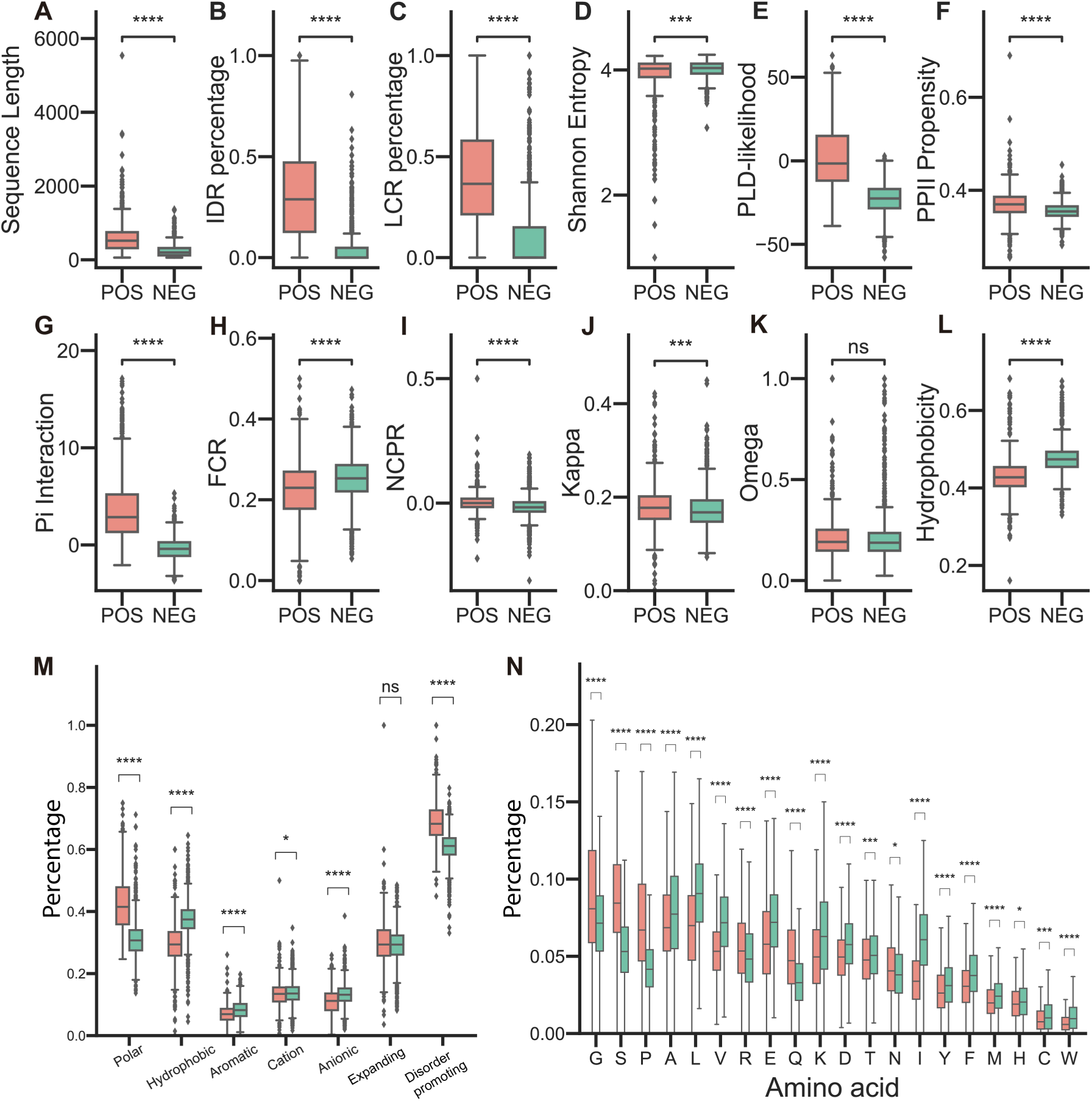
Features Comparison of Phase Separation Positive (POS) and Negative (NEG) Proteins in Training Dataset. The comparison includes (A) Sequence length, (B) IDR percentage, (C) LCR percentage, (D) Shannon entropy, (E) Prion-like domain log likelihood, (F) Type II Polyproline helices propensity, (G) pi-pi interaction, (H) fraction of charged residues, (I) net charge per residue, (J) kappa, (K) omega, and (L) hydrophobicity. Additionally, (M) demonstrates a percentage comparison of different groups of amino acids, and (N) showcases a single amino acid percentage comparison. Features from (H) to (M) are defined by localciders. In all boxplots, the central line illustrates the median value, while the edges of the box denote the 25th (lower) and 75th (upper) percentiles. The whiskers extend to 1.5× the interquartile range from the box edges. Outliers beyond the whiskers’ range are marked as dots. Statistical significance is indicated as *p≤0.05, **p≤0.01, ***p≤0.001, ****p≤0.0001, and ns = not significant (Mann-Whitney test, two-tailed for A-N).

We categorized amino acids into distinct clusters based on their biochemistry properties (Appendix Table 2). Our findings revealed that PS-prone proteins are rich in polar and disorder-promoting amino acids, yet they have fewer aromatic and anionic residues (Fig 1M). Additionally, we observed differences in the composition of amino acids between the POS and NEG datasets (Fig 1N).

### Machine learning model construction and feature evaluation

We selected a total of 39 features to represent the biochemistry properties of proteins involved in molecular condensation (Fig 1). By employing the Uniform Manifold Approximation and Projection (UMAP) algorithm (McInnes *et al*, 2018), we transformed this 39-dimensional data into a 2-dimensional scatter plot. Even without applying a clustering algorithm, a clear distinction between POS and NEG sets is evident, with both forming two separate groups (Fig EV1B). This suggests that the chosen features effectively capture the difference between POS and NEG. To improve the performance of final model, we used the one-sided selection algorithm (Lemaître *et al*, 2017) to undersample the NEG set, ensuring a balanced training dataset. After undersampling, we again used UMAP to visualize the 2D projection, and two sets remain to be distinctly separated (Fig 2A).

**Figure 2.**
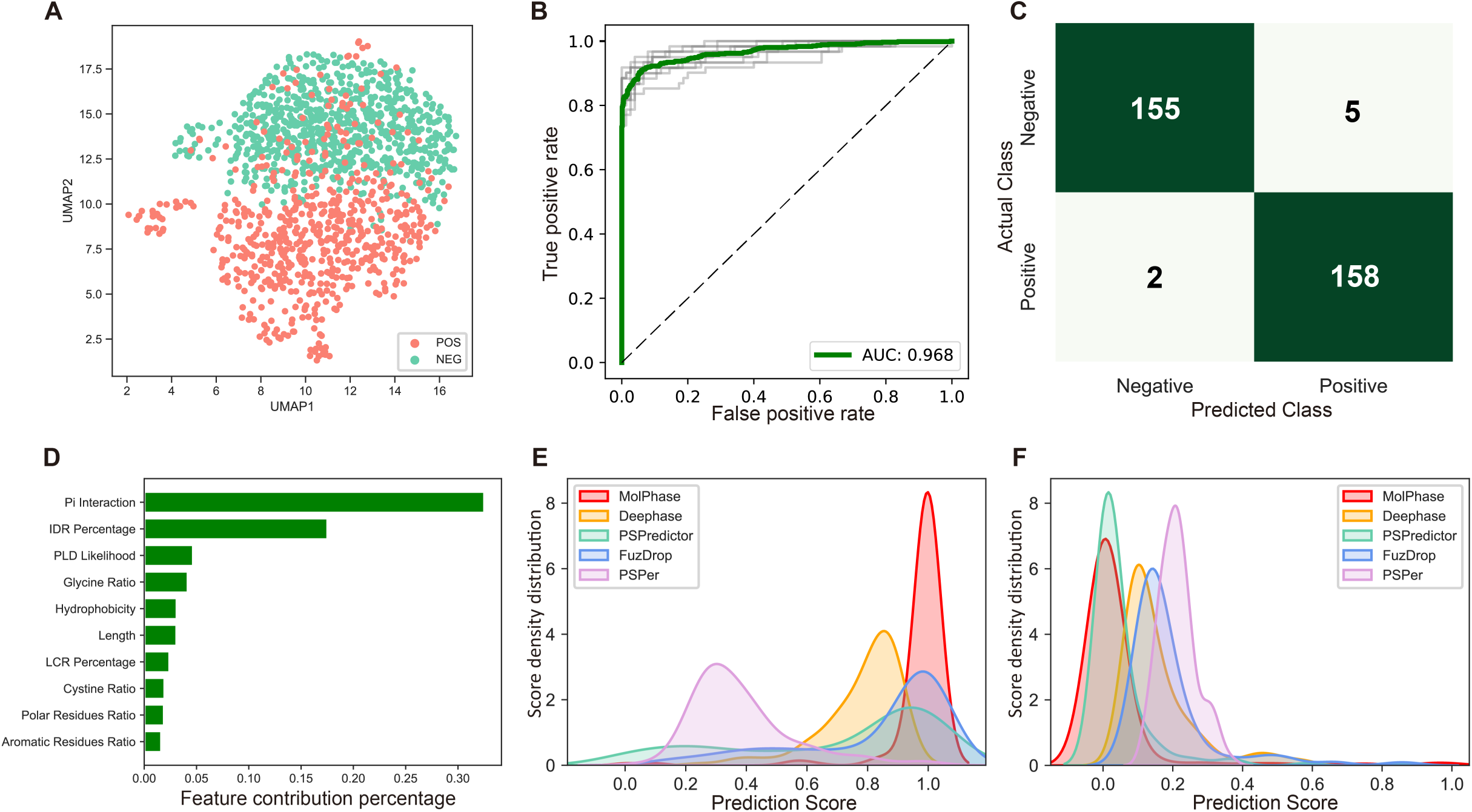
Performance Evaluation of the Machine Learning Predictor, Molecular Condensation and Phase Separation Predictor (MolPhase). (A) UMAP visualization of the 2D vector projection of training datasets following one-sided selection undersampling. (B) Receiver operating characteristic (ROC) curve for the classifier using 10-fold cross validation. The green line depicts the average of all training outcomes, while the gray lines illustrate the ROC curves from 10 training iterations. The area under the curve (AUC) is 0.968. (C) Confusion matrix showcasing predictions for the external dataset made by MolPhase. (D) A representation of the 10 most pivotal features in MolPhase. The x-axis denotes feature importance. (E) Density plot of the prediction probability for the external positive dataset, and (F) negative dataset, as generated by five distinct predictors. On the x-axis is the prediction probability of proteins with LLPS propensity as determined by each predictor, while the y-axis displays the density.

Following the workflow depicted in Fig EV1A, we tested seven different classifier models on the same training set (Fig EV1C). Among these, eXtreme Gradient Boosting (XGBoost) (Chen & Guestrin, 2016) stood out, delivering the best performance (Fig EV1C). Consequently, we selected XGBoost as the final model. We assessed the model’s efficacy using a 10-fold cross-validation and plotted the receiver operating characteristic (ROC) curve. The area under the curve (AUC) was an impressive 0.968 (Fig 2B). For further validation, we employed external datasets to evaluate the performance of our predictor. We utilized the negative testing datasets from DeePhase (Saar *et al*., 2021) and sourced positive and negative PS sequences from PhaSepDB (You *et al*, 2020) and PDB, respectively. To enhance the testing reliability, we excluded sequences from the testing set that bore high similarity to those in the training set. Specifically, we removed the positive testing set with an identity greater than 0.9 and from the negative testing set with an identity above 0.3. The resulting confusion matrix revealed that our model achieved a true positive rate of 96.9% and a true negative rate of 98.7% on the external testing set (Fig 2C). Further analysis of feature contribution uncovered that pi interaction has the most significant effect on molecular condensation (Fig 2D). This is followed by IDR percentage and PLD likelihood. Interestingly, glycine’s ratio accounted for 4.17% in our final model, making it the fourth most important feature. Previous studies have indicated that glycine residues can enhance fluidity in the PS protein(Wang *et al*, 2018). Here, we named our predictor the Molecular Condensation and Phase Separation Predictor (MolPhase).

To evaluate the efficacy of MolPhase, we used the same testing sets to juxtapose its performance with four other previously published predictors: DeePhase (Saar *et al*., 2021), PSPredictor (Chu *et al*., 2022), FuzDrop (Hatos *et al*., 2022) and PSPer (Orlando *et al*., 2019). When compared with these predictors using our positive phase separation protein set, MolPhase displayed the lowest false negative rate (Fig 2C, Fig EV1D-G). Its predicted scores were predominantly clustered around 1 and did not exhibit the pronounced ’long tail’ effect seen in others (Fig 2E). On the negative test set, all five predictors showed similar accuracy rates (Fig 2C, Fig EV1D-G). However, scores from MolPhase and PSPredictor (Chu *et al*., 2022) were predominantly closer to 0 (Fig 2F). In summary, our data underscores MolPhase’s ability to accurately predict both PS positive and negative proteins to provide appropriate weighing in homotypic condensation.

### Application of MolPhase for evaluating phytobacterial effectors’ phase separation in plant host

We sought to validate the MolPhase predictions using a specific biological system. or this purpose, we examined the type III effector (T3E) of pathogenic bacteria both *in vitro* and within host cells. Phytopathogens bacteria utilize the type III secretion system (T3SS) to progressively inject a diverse array of T3E into plant cells throughout the infection process. These T3Es, in a dose-dependent manner, suppress the host’s immune defense, facilitating pathogen’s growth (Jones & Dangl, 2006). Due to the need to navigate through the narrow, needle-like T3SS, T3Es often possess intrinsic properties of high disorder, enhancing their conformational flexibility (LeBlanc *et al*, 2021), which, on the other side, also make them ideal candidates for molecular condensation. We subsequently scrutinized all the T3Es from two extensively researched phytopathogenic model species, *Pseudomonas syringae* pv. *tomato* (*Pst*) DC3000 and *Xanthomonas campestris* pv. *Campestris* (*Xcc*) 8004, based on their annotated genomes(Buell *et al*, 2003; Qian *et al*, 2005). Upon assessing the molecular condensation likelihood of these T3Es, our previously identified phase separation T3E, XopR, emerged as the top contender (Sun *et al*., 2021) (Fig 3A). For further validation, we randomly selected six T3Es - HopS1, HopA1, HopAB2, HopO1-2, XopD2 and XopX2 (Fig 3B and C, Fig EV2A-D) - representing a range of MolPhase propensities. Analyes of their biophysical and biochemical features through MolPhase, combined with structure predictions from AlphaFold2, were shown together (Fig 3B-E, Fig EV2A-D). Notably, while the IDR is important for PS(Gao *et al*, 2022), there’s a stark contrast between the IDR percentages of chosen effectors, HopS1 and HopA1, at 80.5% and 8.9%, respectively (Fig 3B and C), respectively. Moreover, we also incorporated two proteins predicted to be negative, XopQ and mRuby2, for further examination (Fig 3D and E).

**Figure 3:**
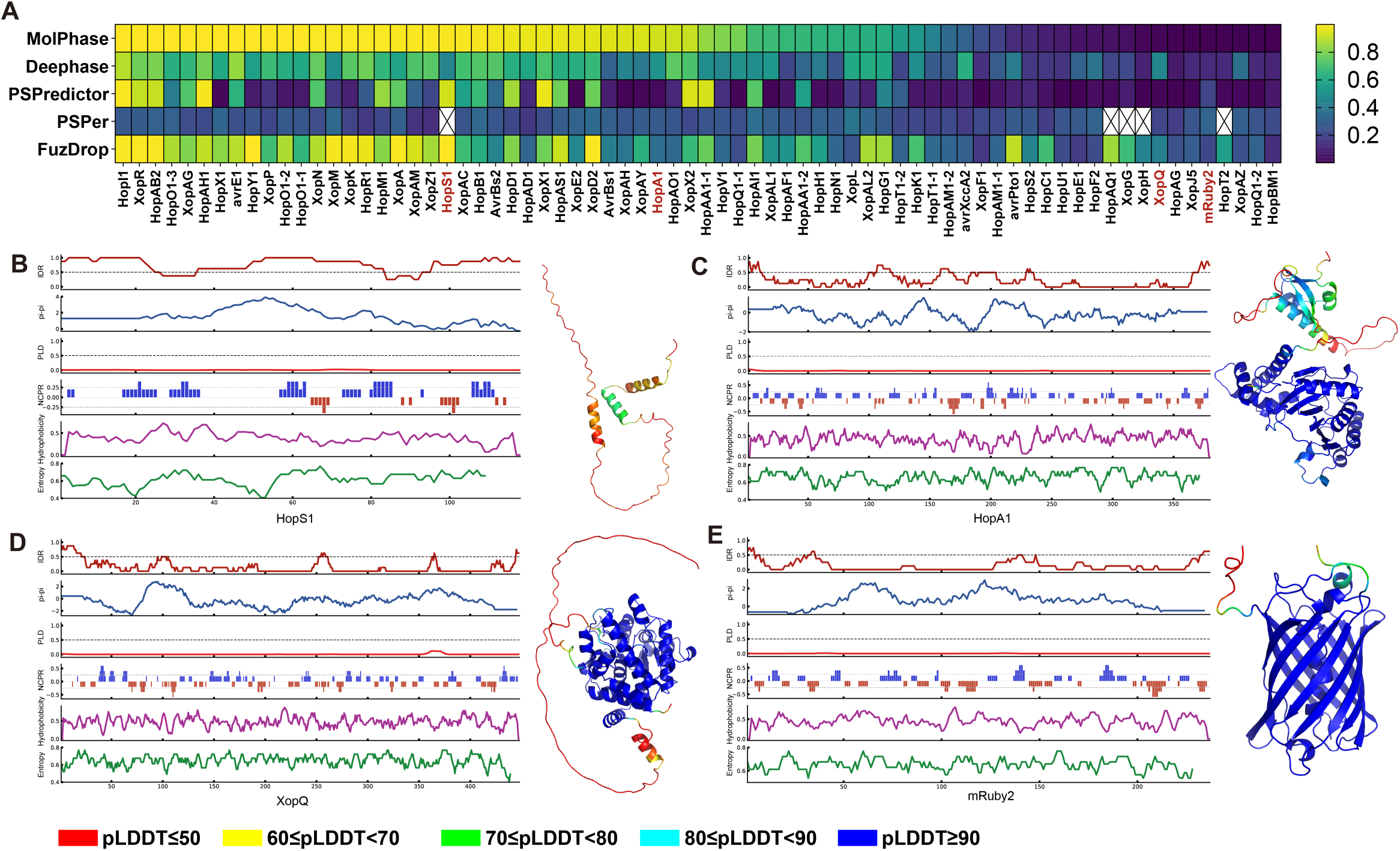
Extraction of Features and Structural Prediction of Effector Proteins. (A) Prediction of phase separation propensity for effector proteins from *Xanthomonas campestris pv. Campestris (Xcc) 8004*, *Pseudomonas syringae pv. Tomato (Pst) DC3000*, and the fluorescent protein mRuby2. Four effectors couldn’t be predicted by PSPer for unspecified reasons and are absent in the heatmap. Effectors are arranged based on the MolPhase score in descending order. Extraction of features and AlphaFold2 structural prediction for two positive candidates: (B) HopS1 and (C) HopA1, and two negative candidates: (D) XopQ and (E) mRuby2. The feature plot sequence is IDR, pi-pi interaction, Prion-like domain likelihood, net charge per residues, hydrophobicity, and Shannon entropy. AlphaFold2 predicted structures are color-coded by the predicted local-distance difference test (pLDDT) show at the bottom of image, with structures having a higher pLDDT indicating higher accuracy. Regions with low pLDDT, especially lower than 50 strongly tend to disorder. The size of the structure images is not to scale.

Next, we proceeded to express and isolate the recombinant proteins of three T3Es and mRuby2 using the *Escherichia coli* system. Securing full-length proteins that are high in intrinsically disordered regions (IDRs) and prone to condensation can be challenging in a few cases due to factors like degradation or precipitation(Alberti *et al*, 2018). However, the full-length HopS1 protein was successfully obtained by co-expressing with its native chaperone, ShcS1, which is located in the same operon as HopS1 (Kabisch *et al*, 2005). The other effector proteins employed in this study were purified using a His-SUMO construct (see Materials and Methods). In vitro, HopS1 and HopA1 exhibited typical phase separation behavior, dependent on protein concentration and ionic strength (Fig 4A and B). Conversely, the MolPhase-predicted negative XopQ failed to form distinct spherical droplets under the conditions tested, forming only sparse tiny punctuates that did not significantly change in size over time (Fig 4C). Another negative candidate, mRuby2, maintained a consistent diffused distribution in solution across all conditions tested (Fig 4D). Notably, while both HopS1 and HopA1 formed spherical droplets, HopS1 displayed higher fluidity than HopA1 in an *in vitro* fluorescence recovery after photobleaching (FRAP) assay. We then assessed their *in vivo* phase behavior by expressing them in *Nicotiana benthamiana* via an agrobacterium-based transient expression system, controlled by an inducible XVE promoter responsive to β-estradiol (Zuo *et al*, 2000). Both HopS1 and HopA1 exhibited two-dimensional condensation on the plasma membrane (PM), in contrast to XopQ and mRuby2, which dispersed throughout the cytosol (Fig 4H). Intriguingly, within plant cells, both HopS1 and HopA1 displayed greater dynamics than their *in vitro* homotypic condensates, as evidenced by FRAP (Fig 4 G and J). This suggests a balanced interplay of inter- and intramolecular interactions *in vivo*, possibly preventing the formation of higher-order and denser assemblies seen *in vitro*, potentially due to the involvement of specific plant proteins or unique intracellular biophysical conditions (Fig EV2A-D). Furthermore, we evaluated four other effectors predicted to be positive *in vivo* in plants. HopO1-2 exhibited 2D phase separation on the PM, whereas HopAB2, XopD2, and XopX2 formed condensates within the cytosol (Fig 4H). In conclusion, all the T3Es examined that were positively predicted by MolPhase displayed phase separation behaviors within live plants upon exogenous expression, mimicking their deposition into the host by T3SS during plant-microbe interactions (Fig 3A).

**Figure 4:**
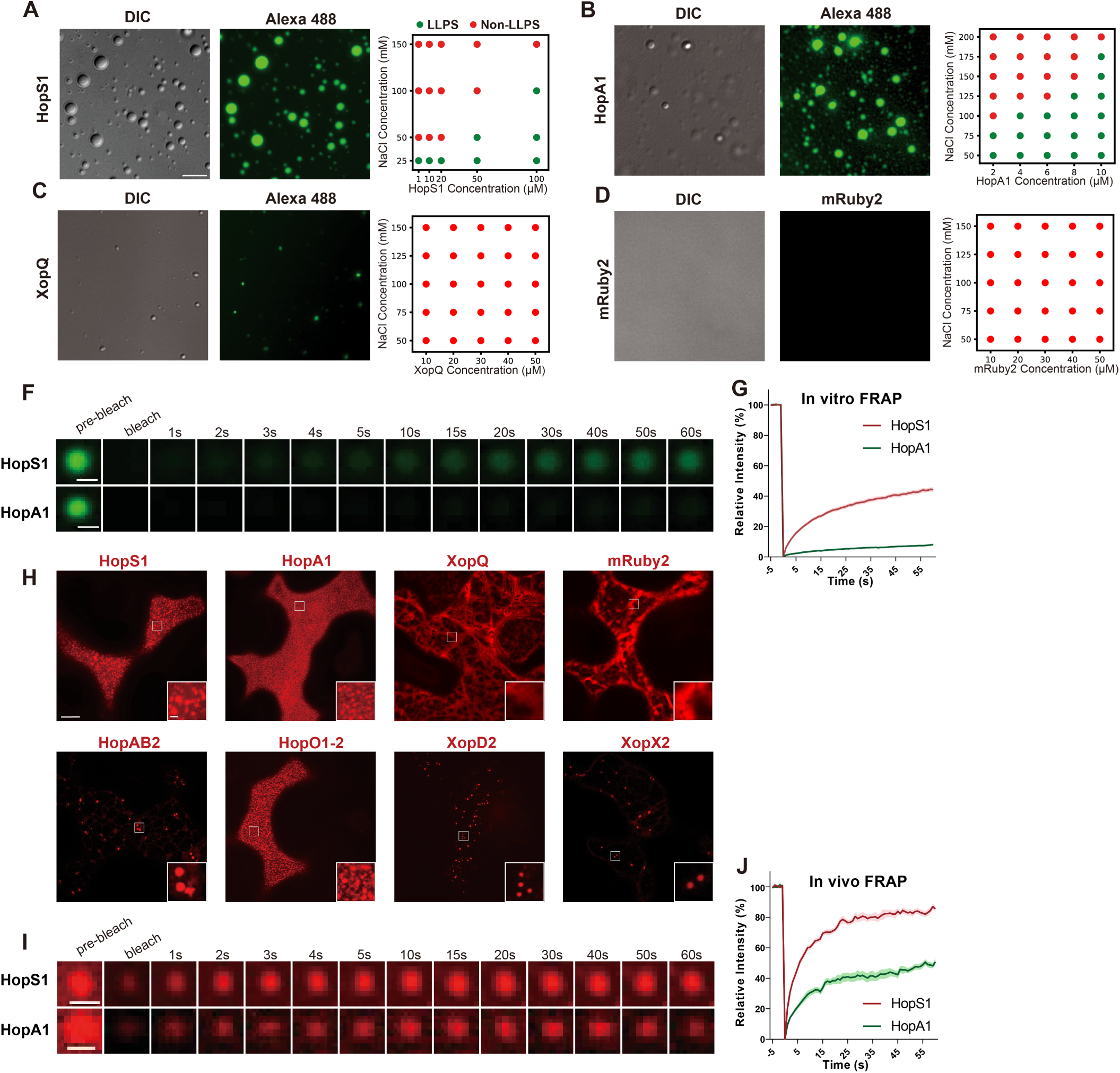
Phase Behavior of Examined Proteins Both *In Vitro* and *In Vivo*. (A)-(D) *In vitro* phase separation studies of proteins predicted for phase separation: (A) HopA1, (B) HopS1, and those predicted against phase separation: (C) XopQ, (D) mRuby2. 1% Alexa 488-labelled protein was combined with 99% unlabelled protein for HopS1, HopA1, and XopQ. Microscope conditions were: (A) 100 μM HopS1 with 25 mM NaCl, (B) 50 μM HopA1 with 50 mM NaCl, (C) 50 μM XopQ with 50 mM NaCl, and (D) 50 μM mRuby2 with 50 mM NaCl. Images have a scale bar of 5 μm. The phase diagram indicates conditions favorable or unfavorable for liquid-liquid phase separation (LLPS). (F) *In vitro* FRAP assay for HopS1 and HopA1 with a scale bar of 1 μm. (G) Relative intensity analysis of the in vitro FRAP assay for HopS1 and HopA1. Intensity prior to bleaching (5s) was averaged and set to 100%, post-bleach intensity was set to 0%, and intensity 1 minute post-bleach was recorded and normalized. (H) *In vivo* tobacco transient expression studies of six proteins predicted for phase separation: HopS1, HopA1, HopAB2, HopO1-2, XopD2, XopX2, and two predicted against it: XopQ and mRuby2. Scale bars are 10 μm for broader images and 1 μm for close-ups. (I) *In vivo* FRAP assay for HopS1 and HopA1. Scale bar = 1 μm. (J) Relative intensity analysis of the *in vivo* FRAP assay for HopS1 and HopA1, using the same intensity calculation methods as in (G). Data in panels (G) and (J) represent 10 independent repeats. The central bold line indicates the mean value, while the shaded areas represent standard deviation.

### Phase separation under proteomic perspective

Taking advantage of MolPhase’s high-throughput capability to predict phase separation (PS) at the proteomics level in living organisms, we analyzed the entire proteomes of *Xcc* 8004 and *Pst* DC3000. Remarkably, the distribution patterns of protein PS in these two strains were quite similar. The majority of proteins exhibited no potential for PS, while only a few proteins demonstrated the ability to undergo PS (Fig 5A and B). We then compared the PS distribution of the phytobacterial proteome with that of their T3Es. To provide a comprehensive evaluation of phytobacterial T3Es, we expanded our analysis to include 529 effectors from 494 strains sourced from *Pseudomonas syringae* Type III Effector Compendium (PsyTEC) (Laflamme *et al*., 2020). As a result, this extensive set of T3Es showed a higher propensity to undergo PS (Fig 5C). The top six features that most influenced the PsyTEC set prediction (Fig 2D, Fig 5D-I) are consistent with MolPhase’s training datasets (Fig 1 B, C, E, G, L and M), suggesting that T3Es have a similar amino acid composition and sequence features to the PS proteins from various species.

**Figure 5:**
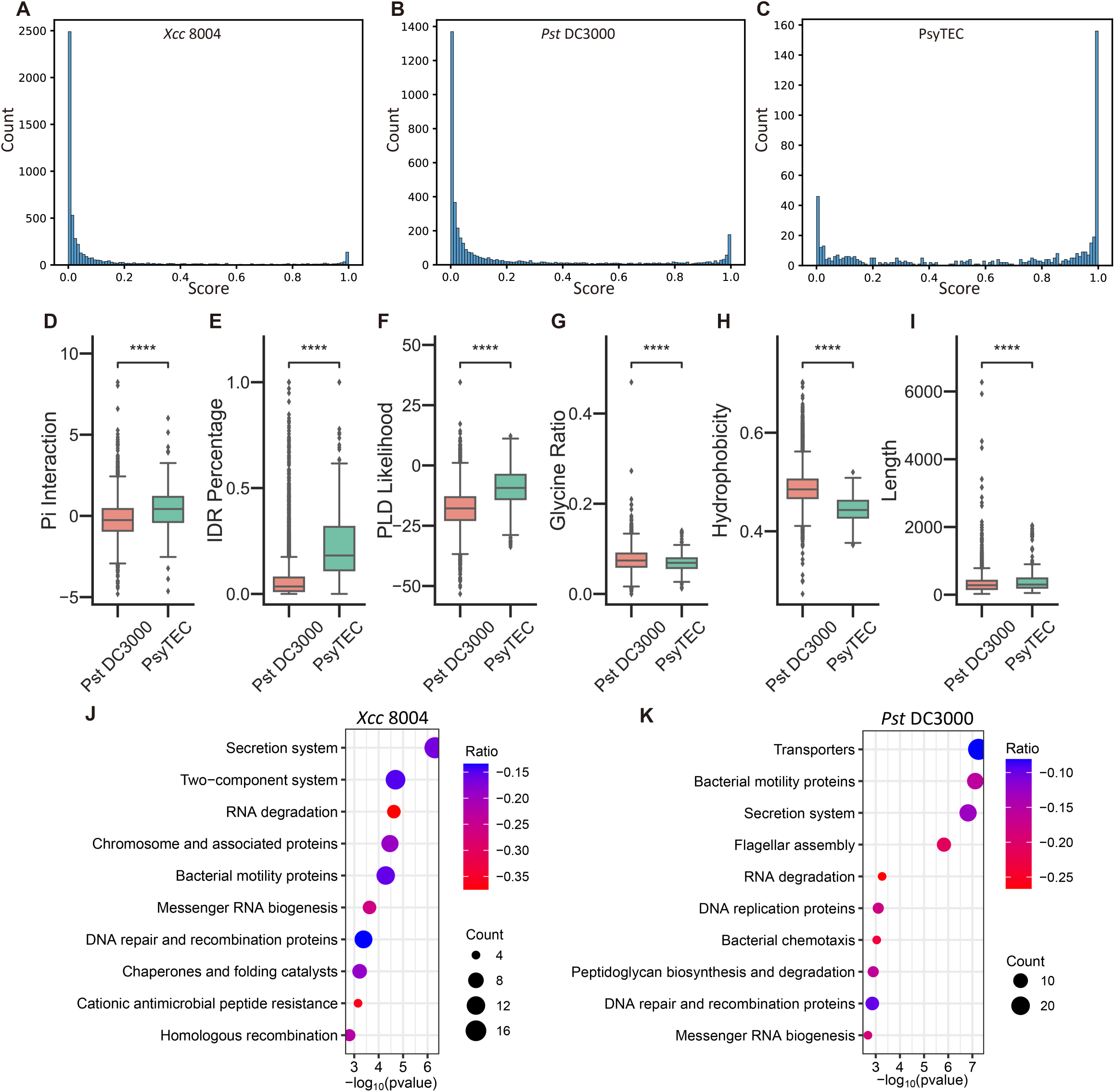
Distribution of Phase Separation Proteins in the Proteome. (A) & (B) Predicted phase separation across the entire proteome for (A) *Xcc 8004* and (B) *Pst DC3000*. (C) Phase separation prediction for 529 *Pseudomonas* effectors from the *Pseudomonas syringae* Type III Effector Compendium (PsyTEC), with the x-axis showing prediction scores in 0.01 increments and the y-axis indicating quantity. (D)-(I) Feature comparisons between the *Pst DC3000* proteome and PsyTEC, including (D) Pi interaction, (E) IDR percentage, (F) PLD-likelihood, (G) Glycine ratio, (H) hydrophobicity, and (I) sequence length. (J) & (K) Kyoto Encyclopedia of Genes and Genomes (KEGG) enrichment analysis for proteins with phase separation prediction scores above 0.9 in (J) *Xcc 8004* and (K) *Pst DC3000*, where the x-axis shows the -log10 p-value for the Fisher exact test, the y-axis lists enrichment items, the ratio indicates the percentage of the entire pathway enriched, and the count shows the number of enriched items. In boxplots from (D)-(I), the central line denotes the median, box edges mark the 25th and 75th percentiles, whiskers extend to 1.5× the interquartile range from the box edges, and outliers are dots. Significance levels are indicated as *p≤0.05, **p≤0.01, ***p≤0.001, ****p≤0.0001, and ns for not significant (Mann-Whitney test, two-tailed in D-I).

To derive functional insights from MolPhase-predicted proteins, we identified proteins with prediction scores above 0.9 and below 0.1 in the proteomes of *Pst* and *Xcc*. were These were designated as phase positive and negative sets, respectively, and then subjected to pathway enrichment analysis (Fig 5 J and K and Fig EV3A and B). Kyoto Encyclopedia of Genes and Genomes (KEGG) enrichment for positive PS proteins revealed several pathways associated with nucleic acids, including RNA degradation, mRNA biogenesis, DNA repair, and recombination (Fig 5 J and K). Intriguingly, these nucleic acid-binding protein candidates are recognized hotspots for PS (Harami *et al*, 2020; Kar *et al*, 2022), aligning with their tendency to form complex condensates with long-chain polyelectrolyte DNA/RNA. This also underscores the potential of DNA/RNA as a versatile interaction platform for their associated proteins. Thus, such interactions, including IDR-IDR associations, could enhance the formation of protein-nucleotide complex PS (Shin & Brangwynne, 2017). Notably, many T3Es from T3SS, categorized under “secretion system”, also showed a propensity for PS in the proteomes of *Xcc* and *Pst.* On the other hand, MolPhase’s predicted PS negative proteins highlighted several consistent pathways in KEGG enrichment, like amino acid metabolisms and ABC transporter. Interestingly, two nucleic acid-related pathways, particularly tRNA biogenesis and ribosome, primarily consisted of predicted PS negative proteins in both *Xcc* and *Pst* proteomes. The precise role of nucleic acids in PS requires more in-depth analysis.

### Online interface for MolPhase

To make MolPhase publicly available and easily accessible, we’ve developed an online interface (http://molphase.sbs.ntu.edu.sg/) (Fig EV4). This platform is constructed using the 39 integrated features (Fig 1). When users input a protein sequence of interest, MolPhase returns a probability score for phase separation (PS) that ranges from 0 to 1. We recommend 0.5 as an initial cutoff to evaluate the queried protein. Furthermore, MolPhase offers a distribution of features along the sequence, assisting users in functional evaluation and hypothesis formulation. This is achieved through a comprehensive sequence scan and a parallel comparison of protein folding and associative motifs, catering to both homotypic and heterotypic interactions. With the insights provided by MolPhase’s detailed analysis and PS predictions, the design of targeted experiments and subsequent functional explorations will be significantly expedited.

## Discussion

### Constructing a higher accuracy phase separation predictor

Biomolecular condensation is primarily driven by a myriad of interaction forces, including hydrophobicity, electrostatic pi/cation-pi, dipole-dipole, and protein-protein/nucleic acid interactions, etc. (Boeynaems *et al*, 2018). Various motifs and regions, equipped with inter- and intramolecular interactions, facilitate the initial association that leads to the formation of higher-order assemblies, such as IDR, PLD, oligomerization motifs, nucleic acid binding motifs etc. Initial generation of phase separation predictors primarily focused on individual interaction forces and incorporated only single or a few features (Vernon & Forman-Kay, 2019), such as PLAAC (Lancaster *et al*, 2014) was based on PLD prediction and PScore (Vernon *et al*., 2018) based on pi-pi contacts. Often an engagement of fewer features lead to suboptimal performance, especially when compared to predictors that integrate multiple features (Chen *et al*., 2022). To offer a more holistic perspective on phase separation, we present MolPhase. It integrates a total of 39 features, making it the most comprehensive tool that addresses the array of interaction forces driving phase separation to date. The superior performance of MolPhase, as depicted in Fig 2E and F, is likely attributed to its ability to factor in more relevant phase separation interactions than other predictors (Chu *et al*., 2022; Hatos *et al*., 2022; Orlando *et al*., 2019; Saar *et al*., 2021). While pi-interactions remain a pivotal factor in MolPhase (Fig 2D), solely relying on pi-pi contacts (Vernon *et al*., 2018) for predicting phase separation often results in reduced accuracy (Chen *et al*., 2022), further emphasizing that phase separation is influenced by various interactions. Narrowing down to only a subset of them might compromise prediction accuracy.

There are now several high-quality databases specifically for phase separation proteins (Hou *et al*., 2023; Li *et al*, 2020; Mészáros *et al*., 2020; Ning *et al*., 2020; Rostam *et al*., 2023). We’ve harnessed the most recent information on phase separation to build our predictor. With curated the experimentally validated homotypic PS protein, we ensured the quality for input data. And we also achieved the largest training set in terms of scaffold PS protein as our known. As these databases continue to grow, we anticipate that the accuracy of phase separation predictors will enhance correspondingly. However, concerning negative training and testing datasets, many predictors, including ours, still rely on sequences from PDB. The PDB mainly catalogues well-structured proteins or domains, often omitting disordered regions that might not undergo phase separation. Thus, utilizing higher quality negative datasets are likely to further improve future PS predictors.

### From MolPhase prediction to condensates properties characterization and potential biological functions

Predicting protein phase separation using sequence information hinges on understanding the biophysical attributes of the protein. This includes its charge, hydrophobicity, and amino acid composition, as they can determine the likelihood of the protein undergoing phase separation. However, the process of protein phase separation is highly dynamic and subject to their surrounding environmental factors, such as the crowding (Delarue *et al*, 2018), concentration of binding partners (Case *et al*., 2019), pH (Adame-Arana *et al*, 2020), temperature (Boeynaems *et al*., 2018), ionic strength (Sun *et al*., 2021). These conditions can modify the interaction and structure of inherent associative motifs, making primary sequence-based predictions of material properties challenging. Nevertheless, juxtaposing a precise MolPhase prediction with structural predictions, such as those made by AlphaFold2 (Jumper *et al*, 2021), can still streamline hypothesis formulation and experimental design for both *in vitro* and *in vivo* functional studies.

Though molecular condensation highly emphasize on the rapid phase behavior switches upon reaching critical concentrations of percolation, phase separation, or density transition (Miao *et al*, 2023; Pappu *et al*., 2023), the biological function also hinges on aspects like spatial distribution, size growth, and molecule partition. These are majorly modulated by nucleation and coalescence over space and time (Lee *et al*., 2023). One key role of PS is to dampen cellular noise by creating concentrated, membraneless compartments (Deviri & Safran, 2021; Klosin *et al*, 2020; Riback & Brangwynne, 2020). To gauge a biomolecule’s noise-canceling potential in various biological settings, MolPhase presents a tailored approach for predicting such behavior. Our validation of MolPhase predictions using phytobacterial T3Es aptly illustrated a plant pathology-related process. Introducing bacterial proteins into plant cells revealed that in vivo PS behavior hinges largely on the inherent attributes of the molecules being examined, without the influence of existing binding partners.

Dissecting the biological functions of phase-separating proteins demands a multidisciplinary approach, intertwining both computational and experimental techniques. This entails localized quantitative molecule measurements, advanced molecular assembly imaging ranging from nanometer to mesoscale, and accurate assessment of functional molecule partitioning and stoichiometry (Case *et al*., 2019; Dine *et al*, 2018; Hubatsch *et al*, 2021; Kent *et al*, 2020; Knerr *et al*, 2023; Lee *et al*., 2023). To design experiments specifically for biomolecules in the context of their physiological or pathological pathways, MolPhase offers an accurate initial screen in a high-throughput manner, allowing for a systematic evaluation by integrating GO annotation, KEGG pathway enrichment analysis, or AlphaFold2-based structural prediction (Jumper *et al*., 2021). Beyond individual proteins, MolPhase can handle whole proteomes, pinpointing pathways abundant with phase separation-prone proteins. This underscores the significance of phase separation in those pathways. This suggests the importance of phase separation in these identified functional pathways. In our comprehensive analysis of prokaryote phytopathogens *Xcc* 8004 and *Pst* DC3000, it emerged that a majority of their proteins don’t undergo phase separation and are predominantly associated with metabolism pathways. This suggests these reactions likely rely on direct binding and precise stoichiometry to deliver quick results, rather than participating in non-linear, equilibrium-based reactions within condensates. On the flip side, the bacterial secretion pathway emerged as a prime candidate for phase-regulated pathways in both strains. In line with this, T3Es analyses of both strains, as well as the *Pseudomonas syringae* Type III Effector Compendium (PsyTEC), revealed a significant number of phase separation-prone candidates. Combined the effectors PS prediction on *Xcc* 8004, *Pst* DC3000 and PsyTEC (Fig 3A and 5C), we shown that phytobacteria effectors are hot candidates for PS (Sun *et al*., 2021). And how they exploit the PS property to subvert plant immunity needs further investigation. By applying MolPhase’s prowess in high-throughput predictions on a wider spectrum, the role of PS underneath evolution could be revealed.

## Materials and Methods

### Reagents and Tools Table

| Reagent/Resource | Reference or Source | Identifier or Catalog Number |
| --- | --- | --- |
| Experimental Models |  |  |
| *Escherichia coli* | N/A | DH5α |
| *Escherichia coli* | N/A | BL21(DE3) Rosetta T1R |
| *Agrobacterium tumefaciens* | N/A | GV3101 |
| *Nicotiana benthamiana* | N/A | N/A |
| **Recombinant DNA** |  |  |
| pER10-XVE-HopS1-mRuby2 | This study | N/A |
| pER10-XVE-HopA1-mRuby2 | This study | N/A |
| pER10-XVE-XopQ-mRuby2 | This study | N/A |
| pER10-XVE- mRuby2 | This study | N/A |
| pER10-XVE-HopAB2-mRuby2 | This study | N/A |
| pER10-XVE-XopD2-mRuby2 | This study | N/A |
| pER10-XVE-HopO1-2-mRuby2 | This study | N/A |
| pER10-XVE-XopX2-mRuby2 | This study | N/A |
| pSUMO-His-HopA1 | This study | N/A |
| pSUMO-His-XopQ | This study | N/A |
| pET-28a(+) ShcS1-HopS1-His | This study | N/A |
| pET-28a(+) mRuby2-His | This study | N/A |
| **Antibodies** |  |  |
| N/A | N/A | N/A |
| **Oligonucleotides and other sequence-based reagents** |  |  |
| PCR primers | This study | Appendix Table 3 |
| **Chemicals, Enzymes and other reagents** |  |  |
| β-estradiol | TCI | E0025 |
| NaCl | Merck | 1.06404 |
| Protease Inhibitor Cocktail Set III, EDTA free | Thermo scientific | A32965 |
| IPTG | Sigma-Aldrich | I6758-5G |
| LB broth | Biobasic | SD7002 |
| MES hydrate | Sigma-Aldrich | M8250 |
| MgCl<sub>2</sub>·6H<sub>2</sub>O | NACALAI TESQUE | 20908 |
| Glycerol | Biobasic | C11056308 |
| Alexa Fluor 488 | Thermo Fisher Scientific | A20000 |
| **Software** |  |  |
| Fiji | https://imagej.net/software/fiji/downloads | Fiji-win64 |
| Prism 9 | https://www.graphpad.com/features | 9.5.0 |
| Python | https://www.python.org/ | 3.8.10 |
| scikit-learn | https://scikit-learn.org/stable/ | 1.1.3 |
| XGBoost | https://github.com/dmlc/xgboost | 1.6.2 |
| UMAP | https://github.com/lmcinnes/umap | 0.5.3 |
| LocalColabFold | https://github.com/YoshitakaMo/localcolabfold | 1.5.1 |
| **Other** |  |  |
| HisTrap HP | GE Healthcare | 17524801 |
| HiPrep Desalting column | GE Healthcare | 17508701 |
| HiLoad Superdex 200 pg preparative SEC columns | GE Healthcare | 28989335 |

### Training and testing dataset construction

We sourced data for positive training dataset (POS) from five expansive LLPS databases: LLPSDB v2.1 (Wang *et al*., 2022a), PhaSePro (Mészáros *et al*., 2020), DrLLPS (Ning *et al*., 2020), PhaSepDB2.1 (Hou *et al*., 2023), CD-CODE (Rostam *et al*., 2023). Additionally, we included experimentally validated phase separation protein sequences handpicked from scientific literature. All data sources are detailed in Appendix Table 1. Filters were set to select protein constructs that can undergo phase separation by themselves, also referred to as scaffold. Since these databases offer different formats and annotations, we manually sifted through the sequences to ensure the reliability of our positive training dataset. Only the experimentally confirmed LLPS scaffold proteins were added to the training set. To eliminate sequences with identity exceeding 90%, we utilized CD-HIT (Fu *et al*., 2012) This resulted in a total of 606 unique sequences for the positive training set. For external validation, we employed a dataset from Saar et al. (Saar *et al*., 2021). This dataset, which comprises 160 sequences recognized for their high phase separation propensity, was drawn from PhaSepDB2.1 (Hou *et al*., 2023).

Our negative dataset, encompassing both training and testing, came from PDB (Berman *et al*., 2000), a repository of structured proteins unlikely to undergo phase separation. We adopted the negative dataset of DeePhase (Saar *et al*., 2021), which contained entirely structured single-chain proteins. After applying a strict 30% similarity cutoff, we randomly chose 1,367 sequences for the negative training dataset, while another 160 sequences comprised the negative testing set.

### Protein features extraction

A comprehensive list of feature extraction methods and associated tools can be found in Appendix Table 2. We standardized all charge calculations to a pH of 7.4. Unless mentioned specifically in Appendix Table 2, default parameters were used. All computational tasks were performed on the Linux Ubuntu 20.04.4 operating system (Canonical Ltd).

### Machine model building

We employed a range of machine learning models, such as support vector machine, decision tree, Gaussian Naïve Bayes, random forest, neural network, and adaptive boosting. These were executed in scikit-learn 1.1.3 (Pedregosa *et al*, 2011) using default parameters. We also used the eXtreme Gradient Boosting (XGBoost) model in xgboost 1.6.2, which offers a speedy and efficient implementation of gradient-boosted decision trees (Chen & Guestrin, 2016). To prune ambiguous and redundant samples from the negative training set, we adopted one-sided selection from Python’s imbalanced-learn 0.9.1 (Lemaître *et al*., 2017). We set n_neighbors to 1 and n_seeds_S to 45. To normalize the input variables, we applied the StandardScaler from Python’s scikit-learn 1.1.3 package to the training dataset.

### Data dimension reduction

The dimension reduction technique was used Uniform Manifold Approximation and Projection (UMAP) based on neighbor graph (McInnes *et al*., 2018). The Python package umap-0.5.3 was empolyed, setting random_state to 42 and n_neighbors to 12.

### Model performance evaluation

During model training and evaluation, we adopted a train-test split ratio of 1:4, supplemented with a 10-fold cross-validation. Moreover, several model performance metrics, including the area under the receiver operating characteristic curve (ROC), accuracy, F1 score, precision and recall, were all calculated using scikit-learn 1.1.3 (Pedregosa *et al*., 2011).

### Protein purification

Genes of HopA1, XopQ, and mRuby2 were cloned into the His-SUMO tag vector and subsequently introduced into *Escherichia coli* BL21 (DE3) Rosetta strain. The bacterial cultures were grown in Terrific Broth until they reached an OD600 of 1. At that point, Isopropyl β-D-1-thiogalactopyranoside was added to a final concentration of 0.5 mM, and the cultures were incubated overnight at 20°C. The cells were then lysed using a microfluidics LM20 microfluidizer. After centrifugation and filtration, the supernatant was loaded onto a 5 mL HisTrap column (GE Healthcare), operated via an ÄKTA™ FPLC system (GE Healthcare). The column was equilibrated with a binding buffer consisting of 20 mM HEPES, 500 mM NaCl, and 20 mM imidazole at pH 7.4. The target proteins were eluted using a gradient increase of the same buffer components but with 500 mM imidazole. The His-SUMO tags were then cleaved off by overnight treatment with SUMO protease at 4°C. Following this, the sample was reloaded onto the HisTrap column to remove the cleaved His-SUMO tag and SUMO protease. Final purification was achieved using a HiLoad 16/600 Superdex 200 pg column (GE Healthcare) with a gel filtration buffer comprising 20 mM HEPES, 500 mM NaCl, and 10% glycerol. HopS1 gene was cloned into pET-28a(+) vector, together with its native chaperone ShcS1 (Kabisch *et al*., 2005). Purification protocol was same as above without removing SUMO tag.

### Protein fluorophore labelling

HopS1, HopA1, and XopQ proteins were labeled using Alexa Fluor™ 488 (Thermo Scientific). The proteins were adjusted to a concentration of 2 mg/mL in approximately 500 mL of gel filtration buffer, then mixed with 0.1 M sodium bicarbonate and 10 µL of Alexa 488 dye (stock concentration of 0.5 mg/mL in DMSO). This mixture was incubated on a rotary shaker overnight at 4°C. Any unbound dye was subsequently removed using a HiPrep™ 26/10 Desalting column (GE Healthcare) connected to an ÄKTA™ FPLC system.

### Nicotiana benthamiana transient expression

*Nicotiana benthamiana* (tobacco) plants were grown around 24℃ with 16-h-light and 8-h-dark cycle. Plasmids carrying the XVE::effector-mRuby2 construct were transformed into the *Agrobacterium tumefaciens* strain GV3010. The agrobacterium cells were cultured in LB medium at 30°C overnight, then harvested by centrifugation and resuspended in a buffer containing 10 mM MES, 10 mM MgCl_2_, and 200 µM acetosyringone. This suspension was incubated for an additional 2 hours at 30°C. Six-week-old tobacco plants were then inoculated using a needle-less syringe to inject the agrobacterium suspension (OD_600_=0.2) into the abaxial leaf surface. After 24 hours, 20 µM of β-estradiol was applied to both sides of the leaves to induce gene expression. Imaging was conducted 24 hours post-induction.

### Microscopy images acquire and fluorescence recovery after photobleaching (FRAP)

For *in vitro* droplet assay, samples were placed on clean cover slides and imaged using a Leica DMi8 (Leica Microsystems) equipped with an HCX PL APO 100x/1.4 oil objective, ORCA-Flash4.0 LT3 sCMOS camera (Hamamatsu), and a solid-state Spectra-X light engine (Lumencor). *In vivo* cell images were acquired on Nikon Ti2 inverted spinning disc confocal (SDC) microscope (Nikon) equipped with a confocal spinning head (Yokogawa CSU-W1), a 100x 1.45NA Plan-Apo objective and a back-illuminated sCMOS camera (Orca-Fusion; Hamamatsu). Excitation light was provided by 488-nm/150mW (Vortran) (for Alexa 488), 561-nm/100mW (Coherent) (for mRuby2) laser combiner (iLAS system; GATACA Systems).

FRAP experiments were performed on the aforementioned Nikon SDC microscope. The laser of 488nm for Alexa 488 and 561nm for mRuby2 were set to 100% power for photobleaching. Timelapse images were acquired with an exposure time of 200 ms and interval of 800 ms in 1 min after photobleaching. All images were acquired using Metamorph software (Molecular Devices) and processed by Fiji.

### Kyoto Encyclopedia of Genes and Genomes (KEGG) enrichment analysis

Proteome sequences of *Xcc* 8004 and *Pst* DC3000 were downloaded from the NCBI (https://www.ncbi.nlm.nih.gov/), KEGG background files were obtained from KEGG GenomeNet (Kanehisa *et al*, 2002). Sequences of interest were send to eggnog-mapper (Cantalapiedra *et al*, 2021) to map respective KEGG pathway k-number. Fisher’s exact test was conducted to calculate the significance of each pathway by Python package, SciPy (Virtanen *et al*, 2020).

## Acknowledgements

This study was supported by MOE Tier 2 (MOE-T2EP30121-0015 and MOE-T2EP30122-0021 to Y.M.), National Research Foundation Singapore NRF-NRFI08-2022-0012; NRF2021-QEP2-03-P10; OF-IRG MOH-000955 that is administered by the Singapore Ministry of Health’s National Medical Research Council, and MOE Tier 3 (MOE2019-T3-1-012) to Y.M. in Singapore.

## Conflict of interest

The authors declare that they have no conflict of interest.

## Author contributions

**Qiyu Liang:** Data curation; formal analysis; investigation; visualization; *in vivo* and *in vitro* protein phase separation assays; methodology; writing – original draft; writing – review and editing. **Nana Peng:** Data curation; formal analysis; investigation; visualization; methodology. **Yi Xie** and **Nivedita Kumar**: protein purification and phase diagram. **Weibo Gao:** Conceptualization; resources; supervision; funding acquisition; investigation; writing – review and editing. **Yansong Miao:** Conceptualization; resources; supervision; funding acquisition; investigation; methodology; writing – original draft; writing – review and editing.

## Data availability

The datasets used in this study are available in the appendix files.

All materials used in this study are available by request from the corresponding author. Source data are provided with this paper.

## Expanded View Figure legends

## Appendix Files Legends

**Appendix Dataset 1: Comprehensive List of Training Set Proteins and Their Respective Data Sources.**

**Appendix Table S1. Definitions and Reference Tools for Features Used in MolPhase Predictions.**

**Appendix Table S2: Classification of The 20 Essential Amino Acids Based on Residue Propensities.**

**Appendix Table S3: Primers Used in This Study.**

**Figure EV1:**
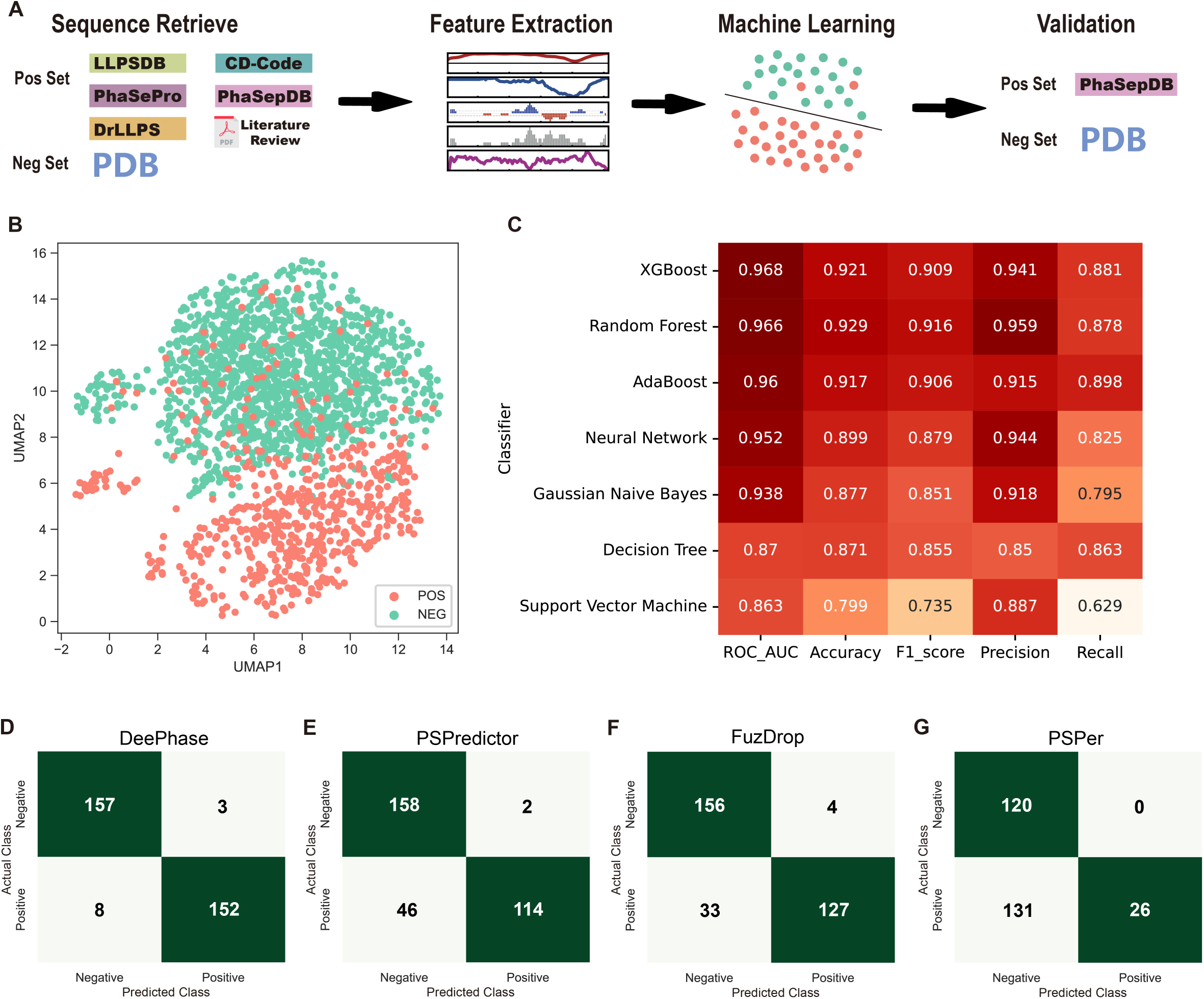
Phase Separation Predictor Performance. (A) Workflow for constructing the phase separation predictor. (B) 2D vector projection of training datasets prior to one-sided selection undersampling by UMAP. (C) Efficacy of seven clustering algorithms on a consistent training set. (D)-(G) Confusion matrices for external dataset predictions by (D) DeePhase, (E) PSPredictor, (F) FuzDrop, and (G) PSPer. FuzDrop’s cut-off is 0.6, while the others are 0.5, as suggested by their respective studies. PSPer couldn’t process 40 sequences for unspecified reasons, which are excluded from the matrix. Deep green indicates true outcomes, while light green indicates false outcomes.

**Figure EV2:**
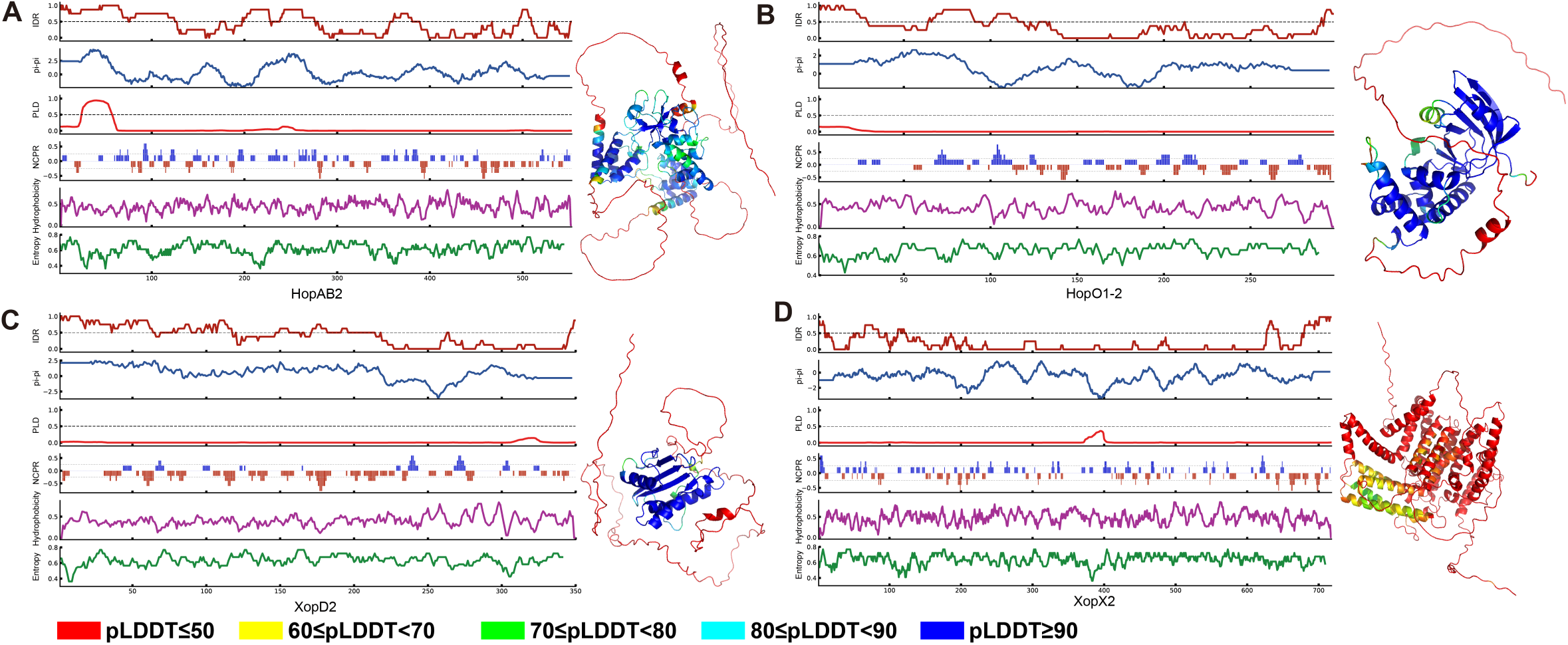
Effector Protein Feature Extraction and Structural Prediction. (A)-(D) Feature extraction and AlphaFold2 structural prediction for four positive candidates: (A) HopAB2, (B) HopO1-2, (C) XopD2, and (D) XopX2. Features, from top to bottom, include IDR, pi-pi interaction, Prion-like domain likelihood, net charge per residues, hydrophobicity, and Shannon entropy. AlphaFold2-predicted structures are color-coded by predicted local-distance difference test (pLDDT) shown at the bottom of image. Structure image sizes aren’t to scale. This relates to Figure 3.

**Figure EV3:**
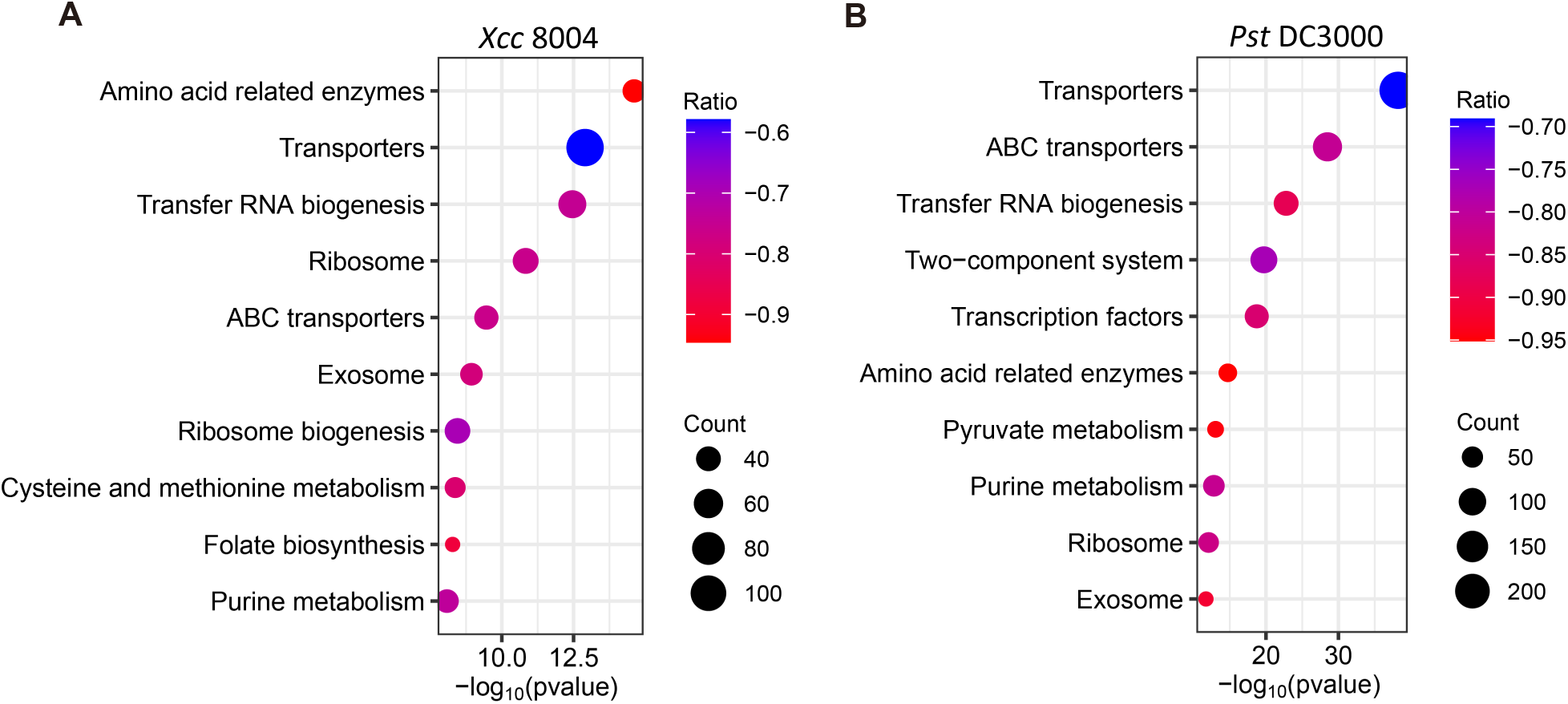
KEGG Analysis for Phase Separation Negative Proteins. (A) & (B) KEGG enrichment analysis for proteins with phase separation prediction scores below 0.1 in (A) *Xcc 8004* and (B) *Pst DC3000*. The x-axis shows the –log_10_ p-value for the Fisher exact test, the y-axis lists enrichment items, the ratio indicates the percentage of the entire pathway enriched, and the count shows the number of enriched items.

**Figure EV4.**
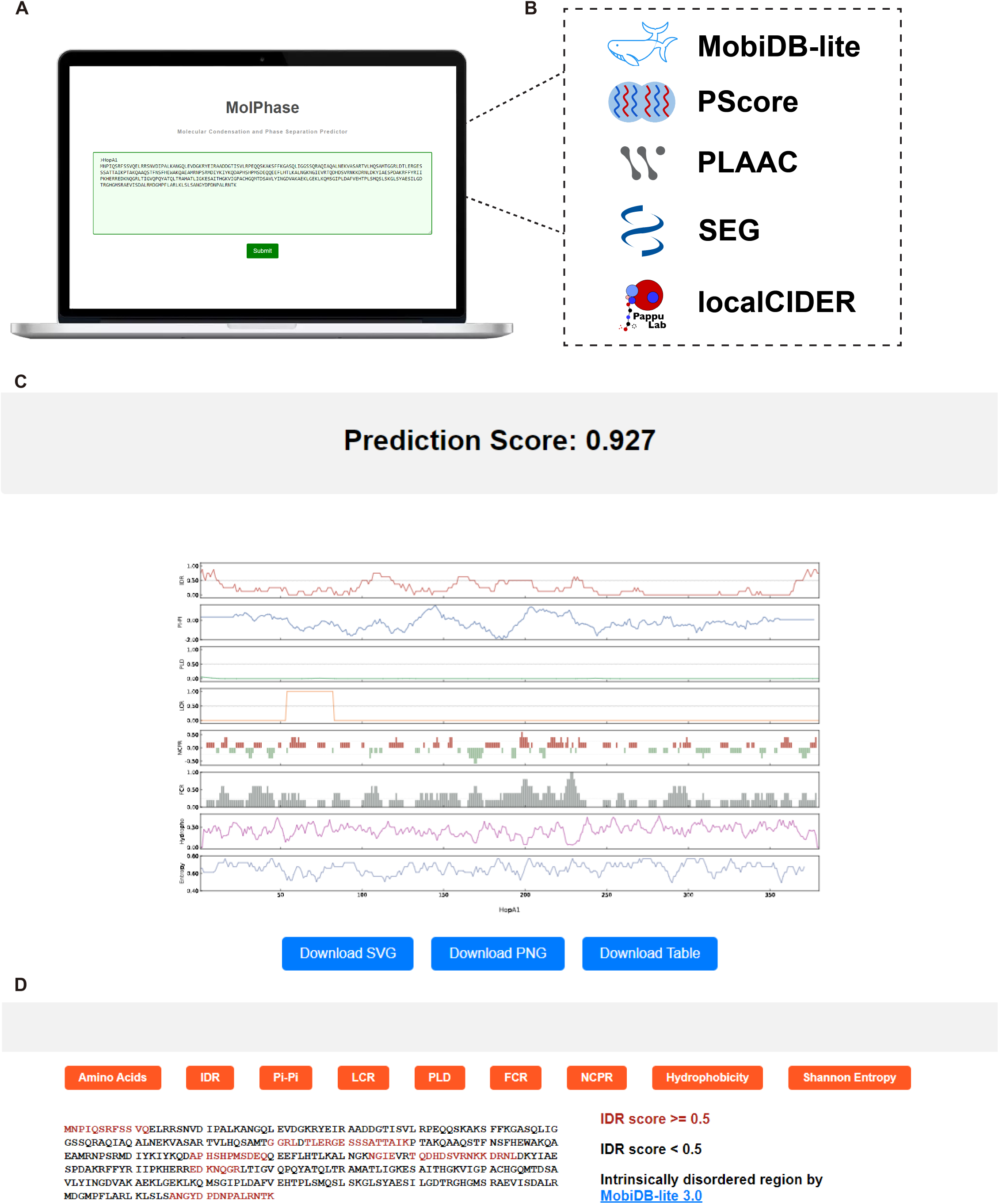
MolPhase Online Predictor Interface. (A) A screenshot of the MolPhase predictor, displaying the input sequence for the effector HopA1. (B) Tools utilized for illustrating features in (C). IDR was determined by Mobidb-lite, pi-pi contacts by PScore, LCR by SEG, and PLD by PLAAC. NCPR, FCR, hydrophobicity, and Shannon Entropy were assessed by localCIDER. The displayed score is the phase separation prediction score, ranging from 0 to 1. (C) Specific features aiding in predicting potential phase separation proteins, using HopA1 as an example. Features, from top to bottom, are IDR, pi-pi, PLD, LCR, NCPR, FCR, hydrophobicity, and Shannon Entropy. (D) Features illustrate in the amino acid sequence view, feature high than threshold score will be highlighted.

